# Sharp wave-ripples in human amygdala and their coordination with hippocampus during NREM sleep

**DOI:** 10.1101/2020.01.07.897413

**Authors:** Roy Cox, Theodor Rüber, Bernhard P Staresina, Juergen Fell

## Abstract

Cooperative interactions between the amygdala and hippocampus are widely regarded as critical for overnight emotional processing of waking experiences, but direct support from the human brain for such a dialog is absent. Using intracranial recordings in four pre-surgical epilepsy patients (two male, two female), we discovered ripples within human amygdala during non-rapid eye movement (NREM) sleep. Like hippocampal ripples, amygdala ripples are strongly associated with sharp waves, are linked to sleep spindles, and tend to co-occur with their hippocampal counterparts. Moreover, sharp waves and ripples are temporally linked across the two brain structures, with amygdala ripples occurring during hippocampal sharp waves and *vice versa*. Combined with further evidence of interregional sharp wave and spindle synchronization, these findings offer a potential physiological substrate for the NREM-sleep-dependent consolidation and regulation of emotional experiences.

## Introduction

Human sleep plays a pivotal role in emotional processing^1^, including the consolidation of emotional memory traces, modulation of emotional reactivity, and regulation of general emotional well-being^2–5^. Such sleep-dependent emotional processing is generally assumed to rely on coordinated activity between the amygdala (AMY) and hippocampus (HPC), as supported by joint AMY-HPC replay of threat-related spiking sequences during animal non-rapid eye movement (NREM) sleep^6^. In contrast, while human neuroimaging studies indicate enhanced AMY-HPC communication during emotional memory retrieval after sleep compared to wake^7^, direct evidence for AMY-HPC communication during human sleep is surprisingly absent.

Ripples, ~80 Hz oscillations found in human HPC^8^ and various neocortical (NC) areas^9–13^, are of potential interest for such AMY-HPC interactions. Sharp wave-ripple complexes in HPC (SPW-ripples; ripples superimposed on ~3 Hz sharp waves), mediate widespread communication between HPC and NC during NREM sleep^14–16^. In animals, neuronal replay preferentially occurs during SPW-ripples^17^, and suppressing SPW-ripples impairs memory consolidation^18^. Importantly, the aforementioned AMY-HPC replay underlying emotional memory consolidation similarly coincides with HPC SPW-ripples^6^, pointing to a key role for ripples in the AMY-HPC dialog. Of note, ripple-like activity has been described in animal AMY^19,20^, raising the possibility of coordinated ripples between these brain structures, but, importantly, ripples have never been described in human AMY.

Beside their close association with SPWs, HPC ripples are nested within HPC and NC ~13 Hz sleep spindles and ~1 Hz slow oscillations (SOs)^8,21^, enhancing HPC-NC information exchange and consolidation^16,22,23^. Whether these additional oscillatory rhythms have a role to play in AMY-HPC communication, either on their own or in conjunction with ripples, also remains unexplored. Here, we report the existence of SPW-ripples in human AMY, and bidirectional AMY-HPC ripple, SPW, and spindle interactions during NREM sleep, offering a potential physiological basis for various forms of sleep-related emotional processing.

## Results

We analyzed invasive electroencephalography (EEG) during NREM sleep from four patients (p1-p4) suffering from intractable epilepsy implanted with multi-contact depth electrodes. We determined bipolar activity from pairs of adjacent contacts located within non-pathological HPC and AMY, as assessed by clinical monitoring and individual anatomy (Fig. 1), ensuring spectral components are generated locally within each brain structure. Sleep architecture calculated from scalp-based polysomnography was in line with normal sleep (Table 1).

**Figure 1.**
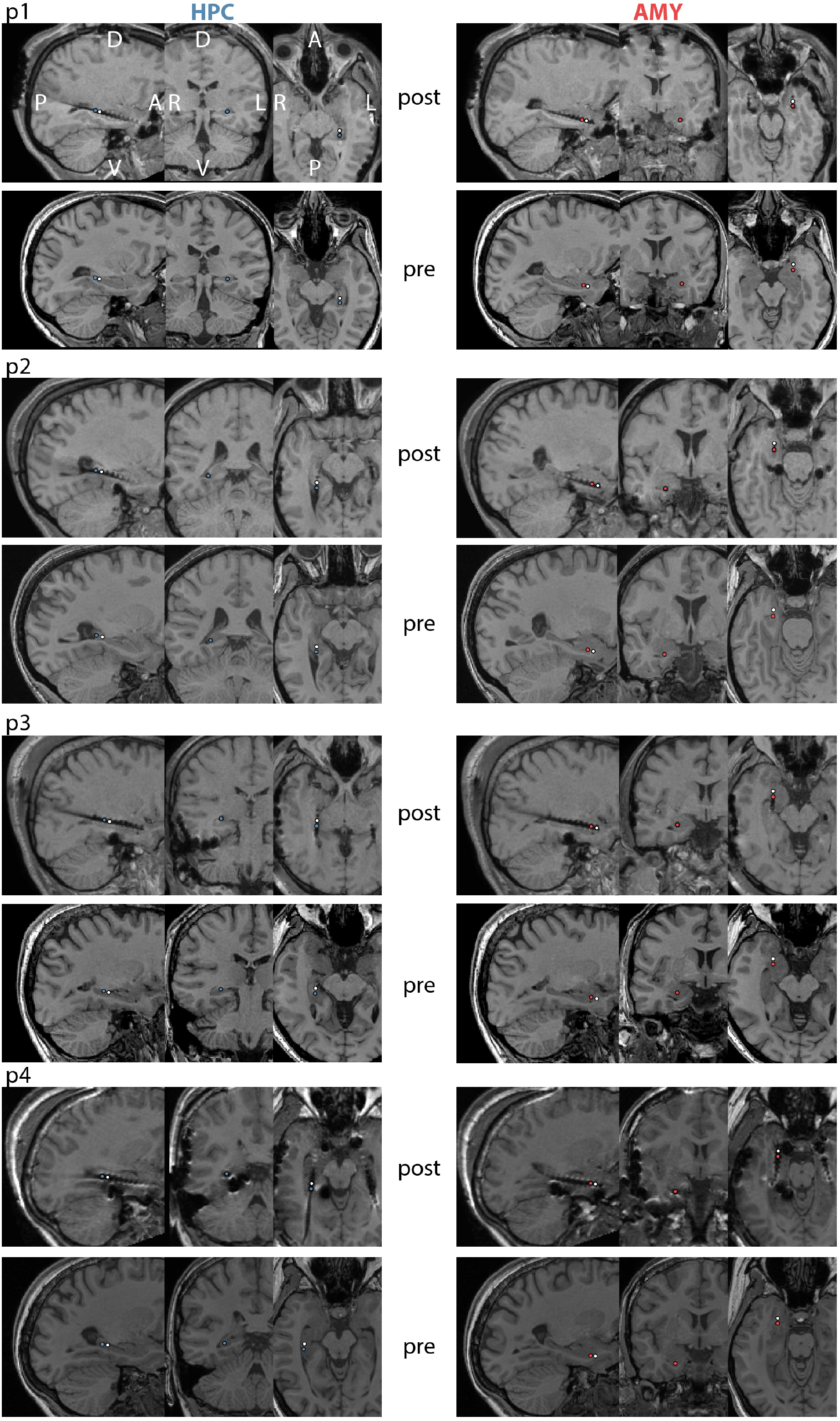
Electrode locations for individual patients. For each patient, top and bottom panels show co-registered post-an pre-implantation T1 weighted MRI scans, respectively. Selected HPC contacts are indicated in left panels (blue: active, white: reference) and selected AMY contacts are shown in right panels (red: active, white: reference). A: anterior, P: posterior, D: dorsal, V: ventral, L: left, R: right.

**Table 1.** Sleep architecture. WASO: wake after sleep onset.

|  | mean | SD |
| --- | --- | --- |
| N1(%) | 27.1 | 15.9 |
| N2 (%) | 45.1 | 10.6 |
| N3 (%) | 11.8 | 7.5 |
| REM (%) | 15.9 | 5.0 |
| N1 (min) | 142.1 | 90.1 |
| N2 (min) | 233.3 | 69.6 |
| N3 (min) | 59.3 | 34.2 |
| REM (min) | 80.2 | 21.8 |
| total sleep (min) | 514.9 | 72.3 |
| WASO (min) | 70.6 | 79.3 |
| sleep efficiency (%) | 88.1 | 12.7 |

### Spectral power and functional connectivity

Group-level spectra adjusted for 1/f scaling showed broad spectral peaks in the 50-100 Hz range for both HPC and AMY (Fig. 2A), comprising the human ripple range^8,21^. Importantly, these ripple peaks were consistently present across patients for both brain structures (Fig. 2B), providing a first indication that ripples may be present in human AMY. Additionally, spindle peaks were present in all four patients for HPC, and in two patients for AMY, consistent with earlier indications of human AMY spindles^24,25^. SO and SPW components were not strongly represented in the adjusted or raw spectra (Fig. 2A, inset) of either brain site, except for an individual with a prominent 4 Hz AMY peak.

**Figure 2.**
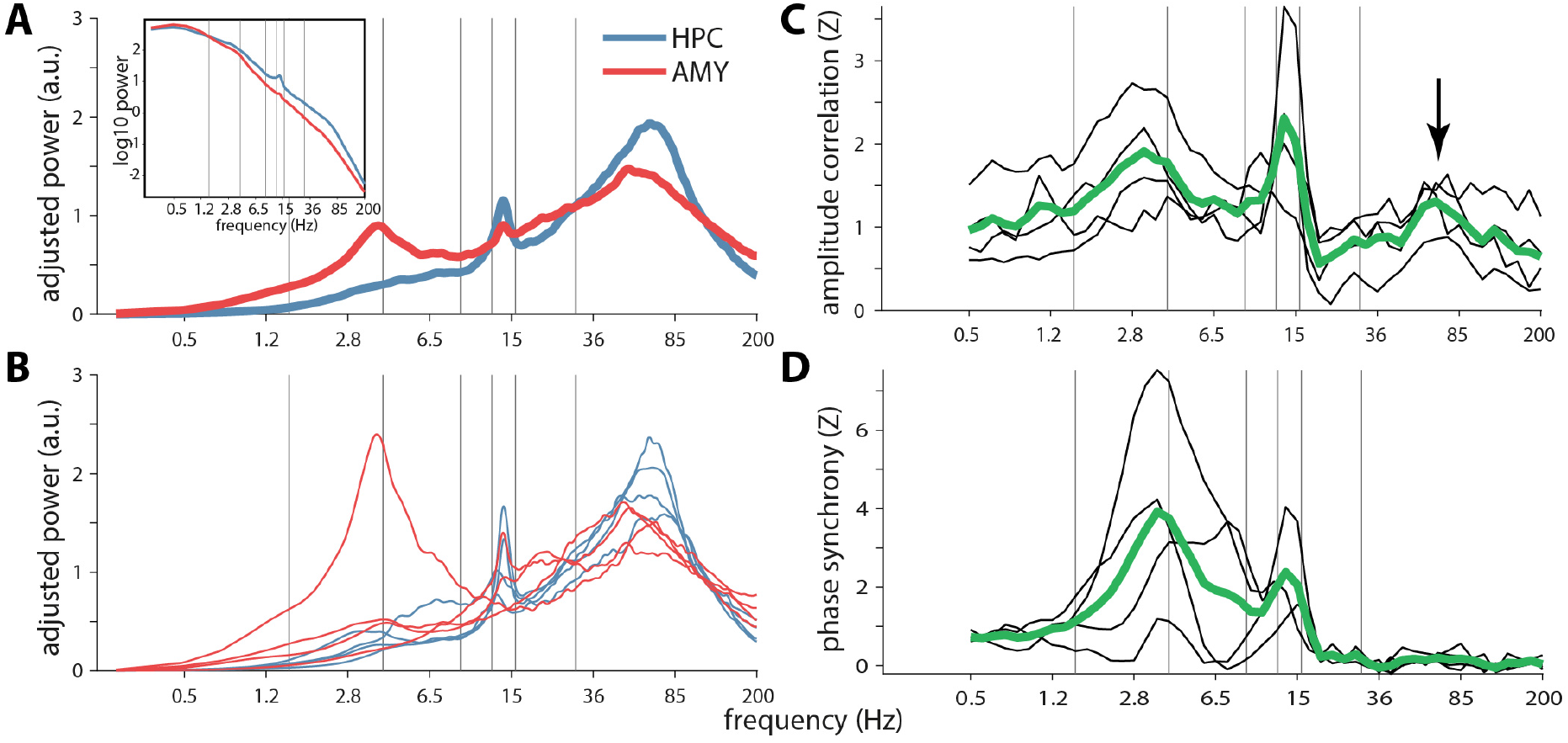
Power and functional connectivity spectra in/between hippocampus and amygdala. (A) Group-level slope-adjusted (main) and raw (inset) spectra. (B). Individual patients’ slope-adjusted spectra. (C) Amplitude envelope correlations between HPC and AMY across patients (green) and for individual patients (black), normalized relative to surrogate distributions. Arrow indicates ripple-band connectivity (D) Phase synchrony, normalized to surrogate distributions. Gray vertical lines at 1.5, 4, 9, 12.5, 16, and 30 Hz indicate approximate boundaries between SO, delta, theta, slow spindle, fast spindle, beta, and faster activity.

Next, we assessed two mathematically and theoretically independent forms of frequency-resolved functional connectivity. Surrogate-normalized AMY-HPC amplitude envelope correlations (Fig. 2C) and phase synchrony (Fig. 2D) signaled robust communication in the SPW (2-6 Hz) and spindle (12-16 Hz) ranges, indicating that activity in these frequency bands both cooccurs and is phase-locked between these brain sites. Importantly, amplitude correlations also peaked in the 50-100 Hz range comprising the ripple band (arrow), suggesting that ripples tend to co-occur between these structures, though with variable phase relations as evident from the lack of phase synchrony.

### Raw traces and spectrograms

Given these initial indications of AMY ripples and their coordination with their HPC counterparts, we visually examined raw HPC and AMY traces along with their spectrograms. This revealed brief (<100 ms) bursts of high-frequency activity in both brain structures centered on the 70-85 Hz range (Fig. 3AB, left). Importantly, many of these events coincided with clear oscillatory behavior in the raw traces (Fig. 3AB, right). Moreover, ripples in both structures were often superimposed on large deflections in the EEG, consistent with SPW-ripple complexes. Interestingly, while HPC and AMY increases in ripple-frequency power were mostly dissociated, instances of co-occurring ripple activity across these brain structures were also observed (Fig. 3B). Moreover, ripples in one site were sometimes associated with SPW-like activity at the other site (e.g., AMY ripple occurring in trough of putative HPC SPW; Fig. 3B). These visual observations, which were similar in the other patients, further suggest the existence of AMY ripples, their association with SPWs, and their coordination with HPC activity.

**Figure 3.**
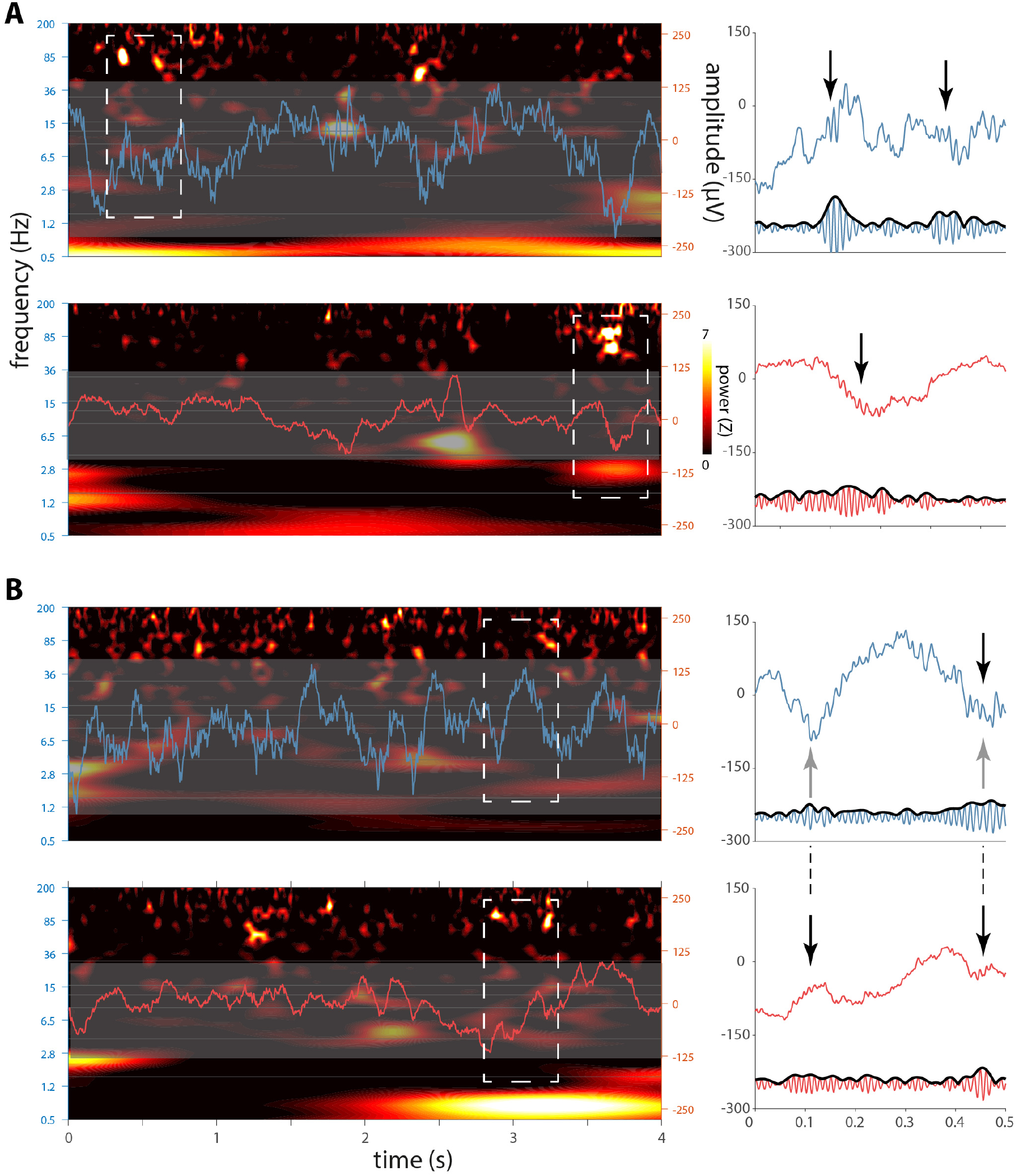
Sleep electrophysiology in hippocampus and amygdala for patient p1. Left panels in (A) and (B) each show 4 s segments of concurrent HPC (top, blue) and AMY (bottom, red) raw traces overlaid on spectrograms (z-scored relative to all NREM sleep). Note brief increases of ~80 Hz power at both brain sites. Close ups of dashed rectangles in right panels indicate ripple-band oscillatory activity in raw (top) and 70-110 Hz filtered (bottom) traces (black arrows: putative ripples), with (A) showing independent HPC and AMY ripples, and (B) showing AMY ripples co-occurring with both a HPC ripple and putative HPC SPWs (gray arrows).

### Ripple characteristics

To examine these possibilities more objectively, we identified ripples using an automated detector (examples for patient p1 in Fig. 4; examples for other patients in Supplementary Fig. 1–3). Across patients, we detected a grand total of 2196 HPC and 979 AMY ripple events. Average ripple density (number per minute) in HPC was consistent with previous human reports^8^ and about twice that of AMY (5.5 ± 1.7 vs. 2.6 ± 1.0; paired t test: t(3)=5.8, P=0.01). Ripple duration (47.9 ± 1.9 vs. 47.1 ± 1.7 ms; t(3)=0.7, P=0.52), main frequency (79.9 ± 1.0 vs. 80.0 ± 1.1 Hz; t(3)=- 0.2, P=0.86), and amplitude (12.1 ± 5.7 vs. 6.3 ± 1.6 μV; t(3)=1.8, P=0.17) did not differ systematically between HPC and AMY, although ripple amplitudes differed between sites on a within-patient basis (Table 2).

**Figure 4.**
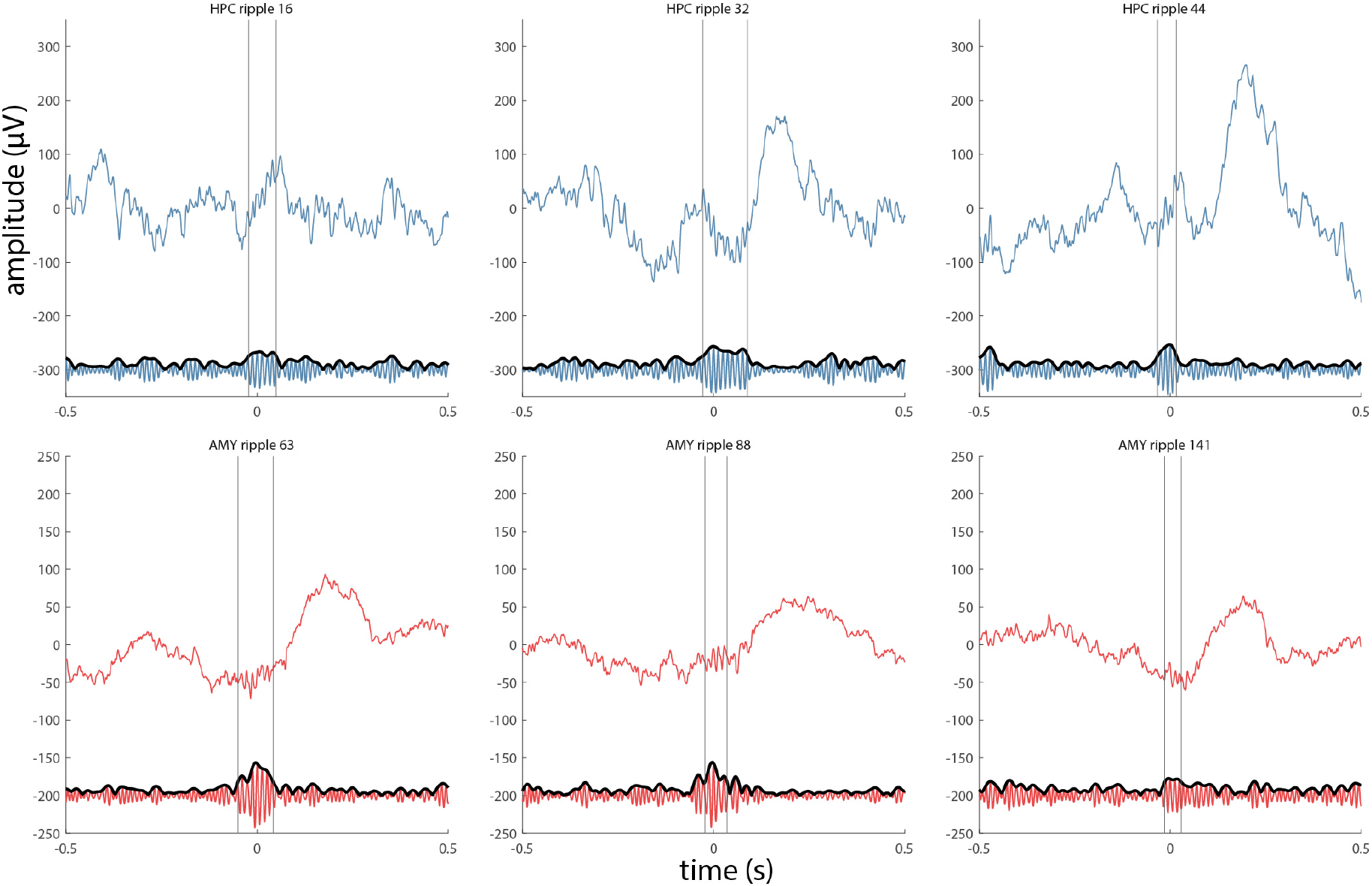
Examples of detected ripples for patient pl. Three HPC ripples (top panels, blue) and three AMY ripples (bottom panels, red) are shown as raw signal (top trace) and in the ripple-filtered (70-110 Hz) band (bottom trace). Vertical lines indicate start and end of ripple.

**Table 2.** Ripple characteristics for individual patients. Results of HPC/AMY comparisons (independent t-test) are shown. Asterisk indicates P value smaller than available numerical precision.

| | | number | density<br>(per min) | duration<br>(ms) | main<br>frequency<br>(Hz) | amplitude<br>(μV) |
| --- | --- | --- | --- | --- | --- | --- |
| p1 | HPC | 353 | 6.2 | $47.6 \pm 11.6$ | $79.6 \pm 6.6$ | $19.3 \pm 3.5$ |
| | AMY | 163 | 2.9 | $44.9 \pm 11.8$ | $81.2 \pm 8.4$ | $6.6 \pm 1.4$ |
|  | *t-test P* |  |  | 0.01 | 0.02 | 0* |
| p2 | HPC | 225 | 3.3 | $45.8 \pm 11.4$ | $79.3 \pm 7.2$ | $12.4 \pm 1.7$ |
| | AMY | 76 | 1.1 | $47.8 \pm 11.6$ | $78.7 \pm 7.9$ | $4.1 \pm 1.1$ |
|  | *t-test P* |  |  | 0.19 | 0.49 | 0* |
| p3 | HPC | 221 | 5.2 | $50.4 \pm 16.6$ | $81.3 \pm 8.4$ | $5.3 \pm 1.3$ |
| | AMY | 138 | 3.3 | $48.8 \pm 12.6$ | $80.6 \pm 8.7$ | $7.7 \pm 2.0$ |
|  | *t-test P* |  |  | 0.32 | 0.46 | 0* |
| p4 | HPC | 1397 | 7.2 | $47.6 \pm 13.1$ | $79.3 \pm 7.7$ | $11.4 \pm 2.0$ |
| | AMY | 602 | 3.1 | $47.1 \pm 12.0$ | $79.5 \pm 8.2$ | $7.0 \pm 1.5$ |
|  | *t-test P* |  |  | 0.42 | 0.56 | 0* |

Next, we calculated the proportion of HPC ripples that co-occurred with the less frequent AMY ripples. Parametrically varying window length, we found relatively low co-occurrence rates of 23.5 ± 6.6 (1,500 ms), 11.2 ± 3.4 (500 ms), and 5.0 ± 1.9 percent (100 ms), indicating that HPC and AMY ripples at the employed recording sites are mostly dissociated. Nonetheless, these cooccurrence rates were significantly enriched relative to surrogate distributions (permutation tests; p1: all P<0.03; p2: all P<0.12; p3: all P<0.003; p4: all P<0.001). As expected, co-occurrences in the opposite direction were lower (11.9 ± 4.4, 5.4 ± 1.9, and 2.4 ± 1.0 percent, respectively), but generally still higher than chance (Table 3). Pooled across all patients’ precise (100 ms window) co-occurrences, ripple timing differences between HPC and AMY did not reliably differ from zero (4.1 ± 28.6 ms; t(63)=1.1, P=0.26). Overall, these findings indicate that a subset of ripples occurs synchronously between HPC and AMY, consistent with the ripple-band envelope correlations of Fig. 2C.

**Table 3.** Ripple co-occurrence rates for individual patients. P values for permutation tests relative to 1000 distributions of surrogate ripples. Asterisk indicates observed ripple co-occurrence was higher than all surrogate iterations.

| window (ms) |  |  | 1500 | 500 | 100 |
| --- | --- | --- | --- | --- | --- |
| p1 | HPC during AMY | % | 25.2 | 11.0 | 4.3 |
|  |  | P | 0.001 | 0.024 | 0.007 |
|  | AMY during HPC | % | 10.5 | 4.5 | 2.0 |
|  |  | P | 0.11 | 0.11 | 0.02 |
| p2 | HPC during AMY | % | 14.5 | 6.6 | 2.6 |
|  |  | P | 0.06 | 0.11 | 0.10 |
|  | AMY during HPC | % | 5.3 | 2.7 | 0.9 |
|  |  | P | 0.07 | 0.06 | 0.11 |
| p3 | HPC during AMY | % | 21.7 | 10.9 | 5.1 |
|  |  | P | 0.003 | 0.004 | <0.001* |
|  | AMY during HPC | % | 16.7 | 7.2 | 3.2 |
|  |  | P | <0.001* | 0.004 | 0.001 |
| p4 | HPC during AMY | % | 32.7 | 16.3 | 8.0 |
|  |  | P | <0.001* | <0.001* | <0.001* |
|  | AMY during HPC | % | 15.1 | 7.2 | 3.4 |
|  |  | P | <0.001* | <0.001* | <0.001* |

### Ripple-related dynamics in HPC and AMY

Next, we examined each patient’s ripple-related dynamics locally within HPC and AMY using five complementary approaches (example patient in Fig. 5AB; other patients in panels AB of Supplementary Fig. 4–6). First, we time-locked the raw signal to the maxima of detected ripples, akin to event-related potential (ERP) analyses (Fig. 5AB, subpanel i). Ripples in both HPC and AMY occurred against a background of large-amplitude fluctuations consistent with SPWs, confirming the individual ripple observations of Fig. 3. These deflections were reliably greater than expected by chance, as indicated by 95% confidence intervals (gray) across 1,000 surrogate ERPs centered on non-ripple events. Importantly, HPC and AMY ripple-related SPW activity was seen for each patient, although precise timing and polarity varied, the latter likely due to bipolar referencing. Second, examination of power spectra for these ripple-centered ERPs revealed strong peaks in the 2-8 Hz SPW range for all patients, with both these SPW peaks and ripple peaks being much more pronounced relative to spectra derived from surrogate ERPs (Fig. 5AB, subpanel ii).

Third, we evaluated ripple-triggered time-frequency power relative to surrogate distributions (Fig. 5AB, subpanel iii). Unsurprisingly, for both HPC and AMY this yielded strong enhancements in ripple-frequency power around the time-locking moment for each patient. Clear increases in 2-8 Hz power surrounding AMY (N=2) and HPC (N=2) ripple detection were also apparent, again consistent with SPW activity. In addition, distinct clusters of spindle power enhancement during or immediately following ripples were seen for AMY (N=2) and HPC (N=1), with an additional patient showing a more broadband power increase comprising the spindle range. Thus, these findings indicate that both SPW and spindle activity tend to occur in close proximity to ripple oscillations, in both AMY and HPC.

**Figure 5.**
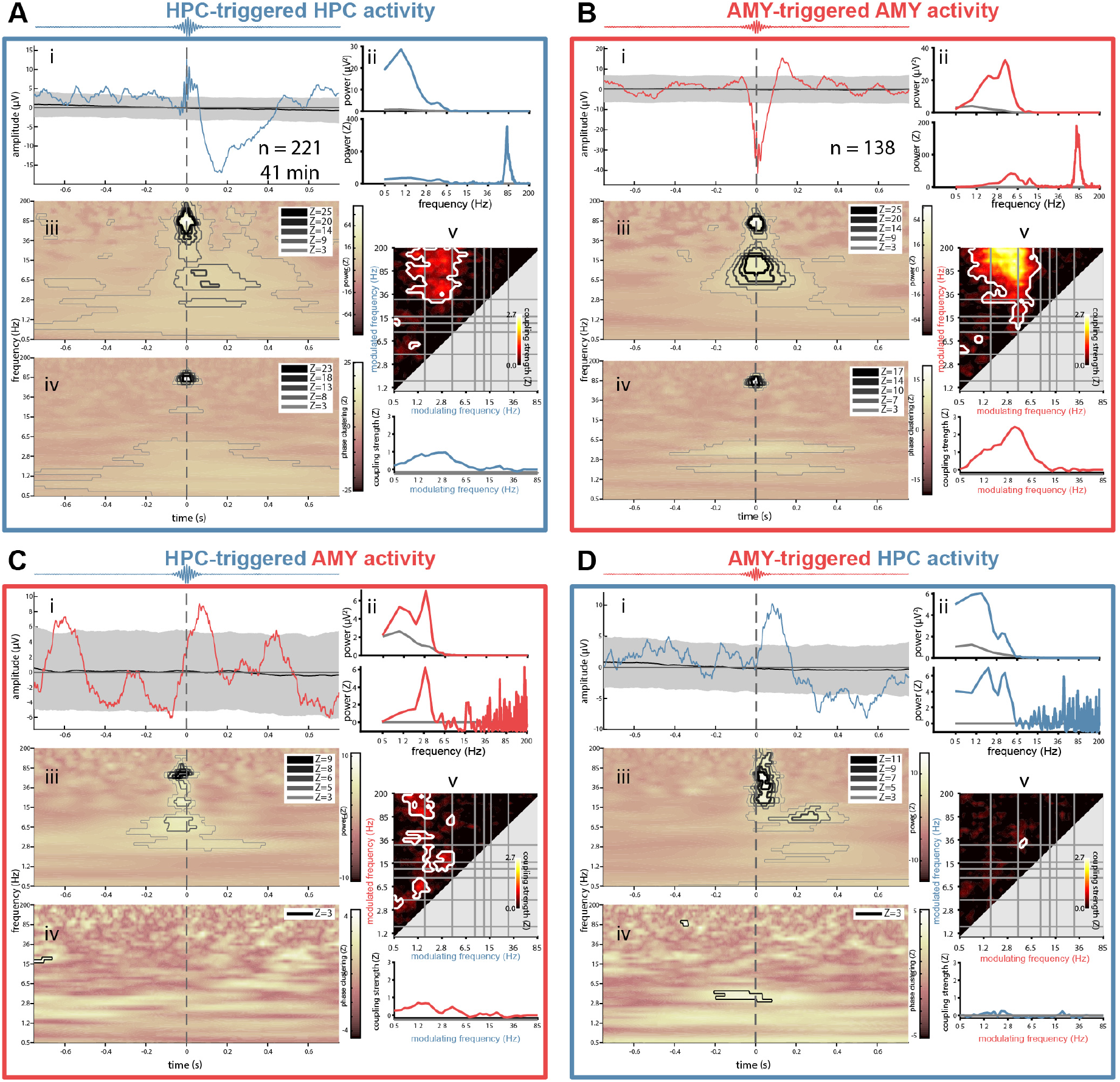
Local and interregional ripple-related dynamics in hippocampus and amygdala for patient p3. Top - local dynamics: HPC activity relative to HPC ripples (A), and AMY activity relative to AMY ripples (B). Bottom – interregional dynamics: AMY activity relative to HPC ripples (C), and HPC activity relative to AMY ripples (D). Subpanels indicate (i): Ripple-triggered ERP (colored) and 95% confidence interval across 1,000 surrogate ERPs, each based on surrogate ripples. (ii): Power spectra of ERPs from (i), with top panel showing raw spectra from ripple ERP (colored) and mean across surrogate ERPs (gray), and bottom panel showing z-scored spectrum (colored) relative to surrogate-based ERP spectra (gray: z=0). SPW peaks are visible in both panels, with additional ripple (A) and (B), and spindle peaks only visible in z-scored spectra. (iii): Ripple-triggered time-frequency power, z-scored relative to surrogates. Contour lines indicate positive (blue) and negative (black/gray) clusters at different levels of significance (Z of 3 and 5 corresponding to P of approximately 0.001 and 10^-7^, respectively). Color scale square root transformed in (A) and (B) to accommodate strong ripple clusters. (iv): Ripple-triggered time-frequency intertrial phase clustering. z-scored relative to surrogates. Clusters as in (iii). (v): Top: comodulogram of cross-frequency phase-amplitude coupling calculated from continuous data, z-scored relative to time-shifted surrogates. For (C) and (D), phase of modulating frequency (x-axis) from “triggered” site (i.e., modulation of AMY activity by HPC phase in (C)). White outlines indicate clusters of significantly higher than zero coupling across 1-min data segments (P < 0.05, cluster-based permutation test). Bottom: modulation of ripple-range (wavelet center frequencies: 75 – 110 Hz) activity by slower frequencies (0.5 - 85 Hz).

The ERP findings from subpanels i suggest that ripples preferentially occur at a specific phase of the SPW. Fourth, therefore, we examined time-frequency-resolved inter-trial phase clustering (ITPC) across ripple trials, relative to surrogate distributions (Fig. 5AB, subpanel iv). Aside from expected cross-trial phase-locking in the ripple band, this analysis indicated consistently phase-aligned activity in the 2-8 Hz range surrounding the ripple maximum for in both HPC (N=4) and AMY (N=4), indicating that ripples are reliably tied to a specific SPW phase. More consistent SPW effects for ITPC than power suggests that some patients’ SPWs do not exceed immediately preceding and following delta/theta-band fluctuations in terms of amplitude, but nevertheless powerfully modulate ripple expression. In addition, we observed clusters of enhanced spindle/beta ITPC around and before the ripple maximum for HPC (N=2) and AMY (N=1), with an additional patient showing larger clusters comprising the spindle range at both sites, further underscoring the relation between ripple and spindle activity.

The preceding indications for ripples being associated with SPWs and spindles are all relative to algorithmically identified ripples, requiring ultimately subjective detection criteria. Previous reports have shown that SPW-ripple activity is also reflected by phase-amplitude coupling (PAC) metrics calculated from continuous data^8,21^, while further allowing the identification of other coupling phenomena. Fifth, therefore, we constructed surrogate-normalized comodulograms from continuous data, indicating the degree of PAC for every frequency pair in the 0.5-200 Hz range (Fig. 5AB, subpanel v). While clusters emerged for various frequency pairs as reported previously for HPC^21^, ripple-band amplitudes in both HPC (N=4) and AMY (N=4) depended strongly on the phase of ~3-6 Hz activity. This is further illustrated by traces at the bottom of each comodulogram, indicating that ripple-band activity is typically coupled most strongly to the delta/theta phase, with additional modulation sometimes exerted by the SO and spindle bands.

Combined, these findings establish that human ripples in both HPC and AMY occur in close temporal proximity to, and are phase coordinated with, local SPWs, and to a lesser extent, sleep spindles.

### Ripple-related dynamics between HPC and AMY

Having characterized ripple-related dynamics locally within HPC and AMY, we turned to cross-regional analyses. Adopting the same analysis strategy as employed in the previous section, we time-locked the raw AMY signal to HPC ripples (Fig. 5C, subpanel i) and the HPC signal to AMY ripples (Fig. 5D, subpanel i; other patients in panels CD of Supplementary Fig. 4–6). This cross-regional analysis again yielded ripple-locked amplitude fluctuations consistent with SPWs in most instances (AMY-locked: N=4; HPC-locked: N=3), albeit with smaller amplitudes (relative to both surrogate ERPs and the local analyses from the previous section). A similar picture emerged from ERP-based power analyses, again expressing clear peaks in the SPW range in most cases (Fig. 5CD, subpanel ii). Importantly, for three out of four patients these effects were present in both directions, suggesting both HPC-AMY and AMY-HPC crosstalk.

Cross-regional ripple-centered time-frequency power analyses (Fig. 5CD, subpanel iii) indicated robust ripple power enhancements at both sites during ripples in the other region (N=2), supporting the earlier ripple-band amplitude correlations and co-occurrence analyses. In contrast, no interregional ripple-band ITPC was seen for these patients (Fig. 5CD, subpanel iv), consistent with the lack of ripple phase synchrony from Fig. 2D. In contrast, for each patient either SPW power or SPW ITPC was enhanced in at least one direction, indicating that ripples at one site are associated with SPWs at the other site. Likewise, interregional comodulogram analyses of continuous data indicated strong bidirectional PAC between the phase of delta/theta frequencies and power in the ripple band (Fig. 5CD, subpanel v), further confirming highly coordinated SPW-ripple activity across HPC and AMY.

## Discussion

While ample behavioral evidence has established a critical role for sleep in emotional processing^1^, it has remained unclear how these processes are implemented neurophysiologically. We report both the existence of SPW-ripples in AMY, and bidirectional electrophysiological AMY-HPC interactions centered on ripple activity during NREM sleep, potentially underlying these behavioral findings.

We demonstrate the presence of ~80 Hz ripple oscillations in human AMY, as indicated by converging evidence from visual examinations, power and functional connectivity spectra, and event detection methods. While this observation is broadly consistent with high-frequency (>120 Hz) ripples in animal AMY^19,20^, and fits with the frequency range of human HPC ripples^8,9,26^, ripples have not yet been reported in human AMY. A first question is whether it is appropriate to employ the term “ripples” for these observations, as SPW-ripples are typically defined in terms of their HPC subfield and laminar generators^27^. On the other hand, ripples have been described in human NC regions both close and distant to HPC^9,10,12,13^. Moreover, given that our detected AMY and HPC ripple events had highly similar spectral compositions and durations, and were similarly associated with local and interregional SPWs, we believe these AMY events may reasonably be categorized as ripples.

HPC SPW-ripples are strongly associated with the replay of task-related firing sequences and subsequent memory consolidation^17,18^. Combined with replay events in AMY^6^, a plausible scenario is that AMY SPW-ripples organize the recapitulation of waking experiences’ affective components. In this light, it is noteworthy that HPC and AMY ripple co-occurrence was modest, suggesting mostly independent reactivation processes at the employed recording sites. However, since ripples can emerge locally anywhere along the HPC axis^28^, many co-occurrences will likely have gone undetected and our co-occurrence rates constitute a lower bound. Still, co-occurrence rates were higher than chance, allowing for integrated AMY-HPC replay, as further supported by ripple-band amplitude correlations (Fig. 2C) and cross-regional ripple power increases for two patients (Fig. 3CD, subpanel iii).

Beside their strong linkage to local SPWs, ripples in AMY and HPC were also reliably associated with SPWs at the other site, expressed as both loose temporal associations and precise phase-amplitude coupling. Similarly, SPW activity was itself coordinated between brain sites, as evidenced by strong enhancements in SPW-band functional connectivity in terms of both amplitude and phase (Fig. 2CD). Overall, these findings indicate that ripples and SPWs, both in isolation and as part of SPW-ripple complexes, are coordinated between HPC and AMY during NREM sleep.

Interestingly, spindles emerged as another oscillatory component mediating AMY-HPC coordination. Irrespective of ripples, both amplitude- and phase-based spindle activity were consistently coordinated across patients between AMY and HPC (Fig. 2CD). These findings extend observations of spindle synchrony between HPC and NC^14,16,29^, or within NC^30^, involving AMY in a widespread network of spindle-related coordination. Moreover, we observed several instances in which ripples were associated with locally enhanced spindle activity in both AMY and HPC. In contrast, evidence for local or interregional spindle-ripple PAC was generally absent (but see Supplementary Fig. 6A, subpanel v). Finally, it deserves mention that no consistent evidence for SO-related AMY-HPC communication emerged, either viewed on its own (Fig. 2CD), or in conjunction with ripples (i.e., no ripple-related SO power or ITPC increases, with one exception in Supplementary Fig. 5C, subpanel iv).

Aside from our choice for local referencing, empirical findings of 1) relatively low ripple co-occurrence, 2) low interregional ripple-band phase synchrony (Fig. 2D), and 3) low interregional ripple-band ITPC (Fig. 5CD, subpanel iv), essentially rule out that AMY SPW-ripples and interregional AMY-HPC communication are due to volume conduction or a common referential signal. At the same time, bipolar referencing required two relatively distant (4.5 mm) contacts to fall inside the small AMY structure, restricting sample size. While this prevented meaningful group-level analyses, evidence for local and interregional SPW-ripples was surprisingly consistent across patients, while linked ripple-spindle activity was also seen in multiple patients. In contrast, various heterogeneous effects (Fig. 5, Supplementary Fig. 4–6) require confirmation in a larger sample.

Patients in our sample exhibited insufficient REM sleep to examine AMY-HPC communication in this brain state, which has also been implicated in emotional (memory) processing^31,32^. However, given the paucity of SPW-ripples^27,33^, replay events^6^, and spindles during REM sleep, any such coordination would likely be implemented differently from the one reported here for NREM sleep. Future work should delineate whether and how ripple characteristics, their co-occurrences, and their linkage to local and interregional SPWs and spindles, are modulated by pre-sleep emotional experiences, or relate to overnight changes in affective (memory) processing.

To conclude, we present first evidence for the existence of AMY SPW-ripples and their coordination with HPC activity. These findings offer an attractive physiological basis for a wealth of findings implicating human sleep, and NREM sleep in particular, in the regulation and consolidation of emotional content.

## Materials and Methods

### Participants

We analyzed archival electrophysiological sleep data in a sample of 4 (2 male) patients suffering from pharmaco-resistant epilepsy (age: 32.8 ± 10.1 yrs, range: 23–47). Patients had been epileptic for 18.8 ± 7.0 yrs (range: 10–27) and were receiving anticonvulsive medication at the moment of recording. All patients gave informed consent, the study was conducted according to the Declaration of Helsinki, and was approved by the ethics committee of the Medical Faculty of the University of Bonn.

### Data acquisition

Electrophysiological monitoring was performed with a combination of depth and subdural strip/grid electrodes. Depth electrodes (AD-Tech, Racine, WI, USA) containing 8–10 cylindrical platinum-iridium contacts (length: 1.6 mm; diameter: 1.3 mm; center-to-center inter-contact distance: 4.5 mm) were stereotactically implanted bilaterally along the longitudinal HPC axis.

Pre- and post-implantation 3D T1-weighted magnetic resonance image (MRI) scans were used to determine electrode locations. Pre-operative T1 (resolution = 0.8×0.8×0.8 mm^3^, TR = 1,660 ms, TE = 2.54 ms, flip angle = 9°) was acquired using a 3.0 Tesla Magnetom Trio (Siemens Healthineers, Erlangen, Germany) with a 32-channel-coil. Post-operative T1 (resolution = 1×1×1 mm^3^, TR = 11.09 ms, TE = 5.02 ms, flip angle = 8°) was conducted using an Achieva 3.0 Tesla Tx system (Philips Healthcare, Best, The Netherlands). Preprocessing and analyses of T1 volumes was done using FMRIB’s Software Library 5.0 (FSL)^34^. Brain extractions^35^ were performed and followed by a bias-field correction^36^. Post-operative volumes were linearly registered to the preoperative volumes. Anatomical labels of the electrodes were determined by an experienced physician (TR) based on these subject-specific co-registered T1 volumes.

For each patient, we selected two pairs of adjacent contacts from the same depth electrode (right: 3, left: 1) contralateral to the epileptogenic side. HPC pairs were located in gray matter (n=3), or on the gray/white matter border (n=1) of the posterior half of the HPC. AMY pairs contained one contact centrally within AMY and one in anterior AMY bordering the temporal pole. For all contact pairs, the more posterior contact was considered the active electrode, and the more anterior one the reference. Distance between HPC and AMY channel pairs was 31.5 ± 3.7 mm (range: 27-36). Additional non-invasive signals were recorded from the scalp (Cz, C3, C4, Oz, A1, A2), the outer canthi of the eyes for electrooculography (EOG), and chin for electromyography (EMG). Signals were sampled at 1 kHz (Stellate GmbH, Munich, Germany) with hardware high- and low-pass filters at 0.01 and 300 Hz respectively, using an average-mastoid reference. Offline sleep scoring was done in 20 s epochs based on scalp EEG, EOG, and EMG signals in accordance with Rechtschaffen and Kales criteria^37^. Stages S3 and S4 were combined into a single N3 stage following the more recent criteria of the American Academy of Sleep Medicine^38^.

### Preprocessing and artifact rejection

All data processing and analysis was performed in Matlab (the Mathworks, Natick, MA), using custom routines and EEGLAB functionality^39^. Preprocessing and artifact rejection details are identical to our previous report^21^. Briefly, mastoid-referenced data were high-pass (0.3 Hz) and notch (50 Hz and harmonics up to 300 Hz) filtered, and channel-specific thresholds (z-score > 6) of signal gradient and high-frequency (>250 Hz) activity were applied to detect and exclude epileptogenic activity. Artifact-free data “trials” of at least 3 s were kept for subsequent processing, resulting in a total of 90.4 ± 70.1 min (range: 42.3-194.3) of NREM sleep.

### Spectral analysis

For each NREM trial and channel, we estimated power spectral density using Welch’s method^40^ with 3 s windows and 80% overlap (0.244 Hz resolution). Mean spectra were determined with a weighted average approach using trial durations as weights. Next, we removed the spectra’s 1/f component to better emphasize narrowband spectral peaks. To this end, we first interpolated the notch-filtered region (50, 100, 150, and 200 Hz, ± 5 Hz) of each spectrum (Modified Akima cubic Hermite algorithm). Then, we fit each spectrum according to *af^b^* using loglog least squares regression^41,42^ and subtracted it from the observed spectrum. Fitting range was restricted to the 4–175 Hz range to avoid the often observed flattening of the spectrum below ~4 Hz and the ~200 Hz notch-interpolated data. Adjusted spectra were resampled to log space and smoothed three times with a moving average window of length 5, as shown in Fig. 2AB.

### Time-frequency decomposition

Continuous data were decomposed with a family of complex Morlet wavelets. Each trial was extended with 5 s on either side to minimize edge artifacts. Wavelets were defined in terms of desired temporal resolution according to:

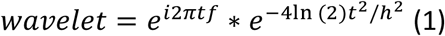

where *i* is the imaginary operator, *t* is time in seconds, *f* is frequency (50 logarithmically spaced frequencies between 0.5 and 200 Hz), *In* is the natural logarithm, and *h* is temporal resolution (full-width at half-maximum; FWHM) in seconds^43^. We set *h* to be logarithmically spaced between 3 s (at 0.5 Hz) and 0.025 s (at 200 Hz), resulting in FWHM spectral resolutions of 0.3 and 35 Hz, respectively. Trial padding was trimmed from the convolution result, which was subsequently downsampled by a factor four to reduce the amount of data. We normalized functional connectivity and PAC metrics using surrogate approaches (see below). To make surrogate distributions independent of variable numbers and durations of trials, we first concatenated the convolution result of all trials of a given sleep stage, and then segmented them into 60 s fragments (discarding the final, incomplete segment).

### Functional connectivity

For every 60 s segment and frequency band, AMY-HPC functional connectivity was assessed using amplitude envelope correlations (AEC)^44^ and the phase locking value (PLV)^45^ as a measure of phase synchrony. AEC was calculated as the Spearman correlation between the magnitudes of the convolution result. PLV operated on the phase angle differences according to:

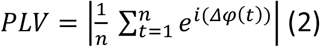

where *i* is the imaginary operator, Δ*φ* indicates phase difference (in radians), and *t* is the sample. We further created normalized version of these metrics using a surrogate approach. Surrogates were constructed by repeatedly (n = 100) time shifting the phase or amplitude time series of one channel by a random amount between 1 and 59 s, and recalculating AEC and PLV for each iteration. These distributions were then used to z-score raw AEC and PLV values, as used in Fig. 2C and 2D, respectively.

### Cross-frequency phase-amplitude coupling

For every 60 s segment, PAC was determined between all pairs of modulating frequency *f1* and modulated frequency *f2*, where *f2*>2\**f1*. We employed an adaptation of the mean vector length method^46^ that adjusts for possible bias stemming from non-sinusoidal shapes of *f1*^47^. Specifically, debiased phase-amplitude coupling strength (dPAC) was calculated as:

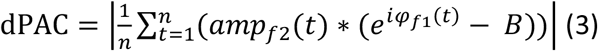

where *i* is the imaginary operator, *t* is time, *amp_f2_(t)* is the magnitude of the convolution result, or amplitude, of *f2*, *φ_f1_ (t)* is the phase of *f1*, and *B* is the mean phase bias:

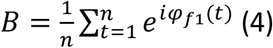

For same-site PAC (i.e., within HPC or within AMY) *φ_f1_* and *amp_f2_* stemmed from the same channel, whereas cross-site PAC used phase information from one brain structure and amplitude information from the other. For every 60 s segment, frequency pair, and same/cross-site combination we constructed a surrogate distribution of coupling strengths by repeatedly (n = 100) time shifting the *f1* phase time series with respect to the *f2* amplitude time series, and recalculating the mean vector length for each iteration. We then z-scored the observed coupling strength with respect to this null distribution of coupling strength values to obtain dPAC_z_, as shown in subpanels v of Fig. 3.

### Ripple detection and surrogates

NREM channel data was zero-phase band-pass filtered between 70 and 110 Hz with 5 Hz transition zones. The ripple envelope was calculated as the magnitude of the Hilbert-transformed filtered signal. Whenever the z-scored envelope exceeded an upper threshold of 2.5 a potential ripple was detected, while crossings of a lower threshold of 2 before and after this point marked the beginning and end, respectively, of the ripple. Start and end points were required to be at least 35 ms apart, corresponding to approximately three full cycles at 70 Hz. Ripple events that did not contain a minimum of 0.75 s of clean data on either side (corresponding to 1.5 s window lengths for co-occurrence and time-locking analyses) were discarded. Duration, maximum amplitude of ripple-filtered signal, and frequency of each ripple were determined, as was ripple density (number per minute).

For each patient and channel, we constructed 1,000 distributions of surrogate ripples, with each distribution containing as many surrogate ripples as detected ripples. Specifically, each surrogate ripple was defined as a random time point within the NREM record, provided that this time point had a minimum of 0.75 s of clean data on either side, and that this extended 1.5 s window did not overlap with a true ripple’s 1.5 s window. Note that while this approach allows overlapping data windows between surrogate ripples, the exact samples used for surrogate cooccurrence and time-locking analyses will seldomly overlap.

### Ripple co-occurrence

For both AMY and HPC channels, and for each detected ripple, we counted a cooccurrence when that ripple’s maximum occurred within an interval (1.5 s, 500 ms, or 100 ms) surrounding any ripple maximum in the other channel. Surrogate co-occurrence rates were determined between a channel’s true ripples and each of the 1,000 surrogate ripple distributions from the other channel.

### Ripple-locked analyses

All local and interregional ripple-related dynamics considered a 1.5 s analysis window centered on the ripple maximum, and the 1,000 distributions of surrogate ripples. Event-related potentials (subpanels i of Fig. 3) were determined by averaging mean-centered ripple trials. The same procedure was used for each surrogate distribution, and the resulting distribution of surrogate ERPs was used to determine the 95% confidence interval at each time point. True and surrogate ERPs were further subjected to spectral analysis (Welch: window length 1.25 s, overlap 95%, spectral resolution 0.488 Hz), and ripple-related ERP power was visualized in raw and surrogate-normalized formats (subpanels ii of Fig. 3).

Time-frequency power (squared magnitude of convolution result) was first normalized (z-scored) relative to all NREM sleep, followed by averaging across ripple-centered trials. This procedure was repeated for each surrogate distribution, and the true mean ripple-related power response was z-scored relative to the 1,000 mean surrogate responses (subpanels iii of Fig. 3). Time-frequency resolved ITPC (subpanels iv of Fig. 3) was determined similarly, both across true ripple trials and each distribution of surrogate trials, followed by z-scoring. Calculations followed equation (2), but using absolute phases instead of phase differences, and averaging across trials rather than time.

### Experimental design and statistical analyses

With the exception of group-level comparisons of ripple properties (paired t test), statistical analyses were performed at the individual level. Significance of ripple co-occurrence was assessed by determining the proportion of surrogate co-occurrence rates that were identical or larger than the observed co-occurrence rate. Reliability of ripple-locked ERPs may be assessed by comparing their amplitude to the 95% confidence interval across surrogate ERPs. ERP-based power spectra are shown both in raw format and z-scored relative to surrogate ERP spectra, with a one-to-one mapping between z-scores and P values (e.g., z-scores of 2, 3, and 5 correspond to one-sided P values of approximately 0.02, 0.001, and 10^-7^, respectively).

Ripple-locked power and ITPC responses, normalized to surrogate distributions, often yielded very high z-scores (particularly for local ripple-band responses). Hence, rather than evaluating significance at a single statistical threshold, cluster outlines were determined at a maximum of five integer z-values, ranging between the lowest Z≥3 not resulting in a cluster comprising all frequency bands, and the highest Z≤25 still generating a cluster. Clusters were required to span at least two frequency bins and ten time bins (36 ms). Note that Z=5 corresponds to strict Bonferroni correction for a two-sided test across all time-frequency points (0.025/18,850≈10^-6^).

The presence of PAC was assessed using cluster-based permutation tests^48^ with a *clusteralpha* value of 0.1 and 1000 random permutations. Specifically, dPAC_z_ values at each frequency pair were compared to zero across data segments using one-tailed t tests (only above-zero effects are of interest). Clusters were required to span at least 2 x 2 frequency bins, and were deemed significant at P<0.05 (one-tailed).

Of note, variable amounts of continuous data, and variable numbers of detected ripples, imply differential statistical power. Consequently, patients/channels with more data tend to show stronger effects (e.g., larger z-scores and cluster extents for p4). Although limiting analyses to identical amounts of data across patients addresses this issue, we did not wish to discard valuable data (e.g., 95% of p4’s HPC ripples should be removed to match p2’s number of AMY ripples), particularly in light of our small sample. It is partly for this reason that we present data on an individual basis with variable data-derived statistical thresholds.

### Data and code availability

Data are not publicly available due to privacy concerns related to clinical data, but data and accompanying analysis code are available from the corresponding or senior author upon obtaining ethical approval.

## Acknowledgments

This work was supported by the German Research Foundation (FE366/9-1 to JF) and a Wellcome Trust/Royal Society Sir Henry Dale Fellowship (107672/Z/15/Z to BPS).

## Author contributions

Conceptualization by RC and JF; methodology by RC, TR, and JF; analysis by RC; interpretation by RC, BPS, and JF; visualization by RC and TR; writing by RC and JF; funding acquisition by JF.

## Competing interests

The authors declare no competing interests.

## Supplementary Figures

**Supplementary Figure 1.**
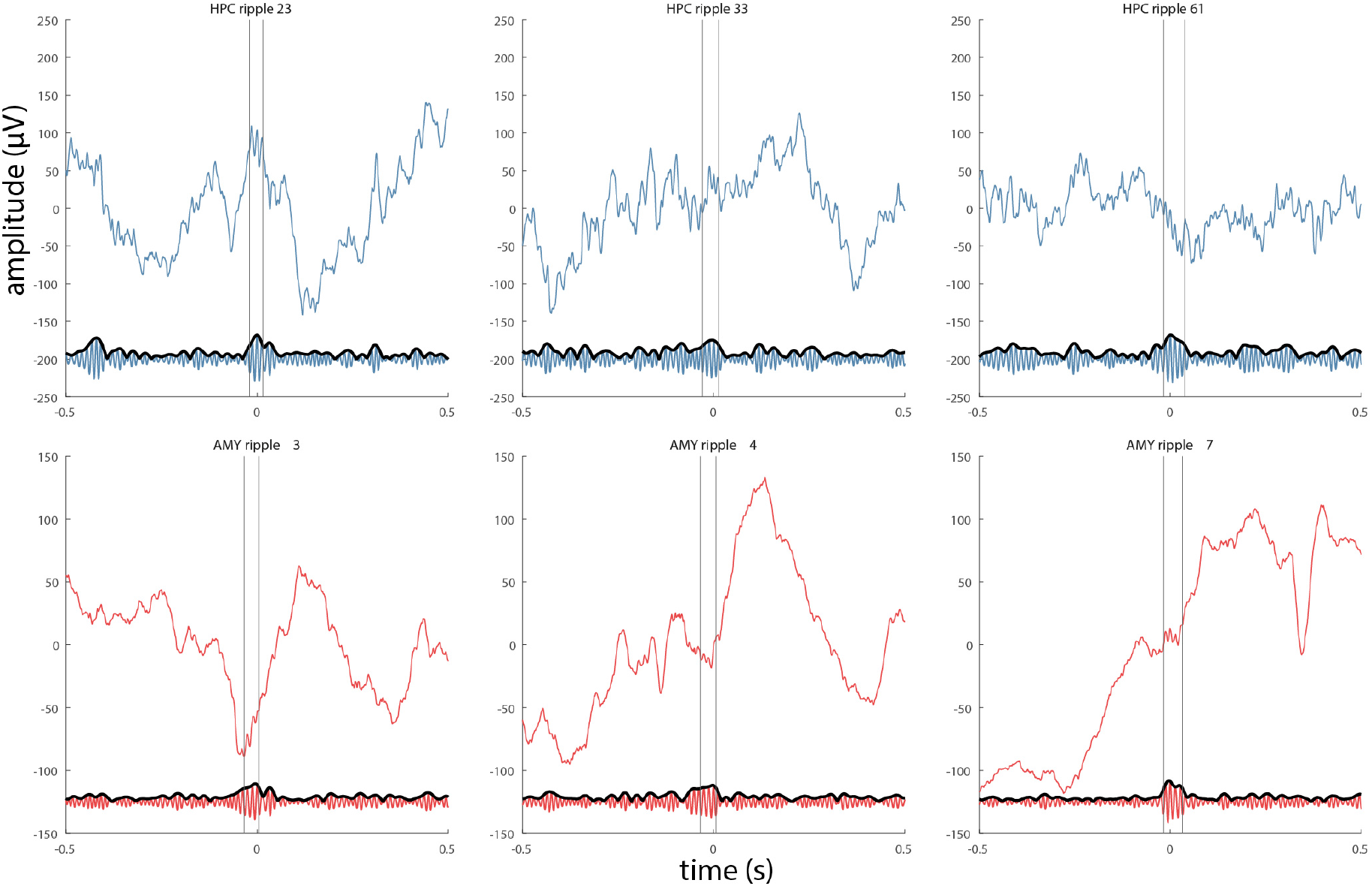
Examples of detected ripples for patient p2. Figure layout as in Figure 4.

**Supplementary Figure 2.**
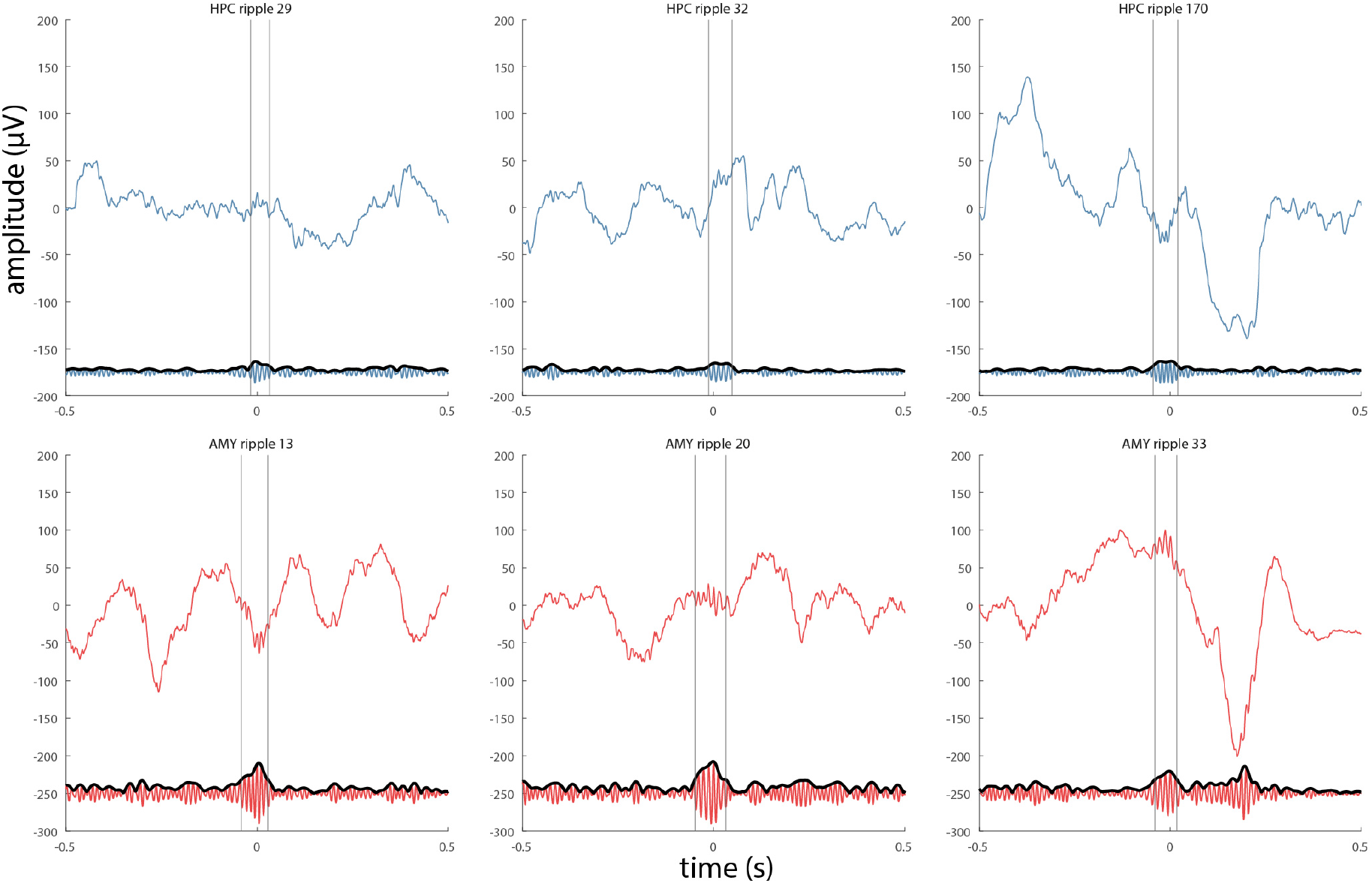
Examples of detected ripples for patient p3. Figure layout as in Figure 4.

**Supplementary Figure 3.**
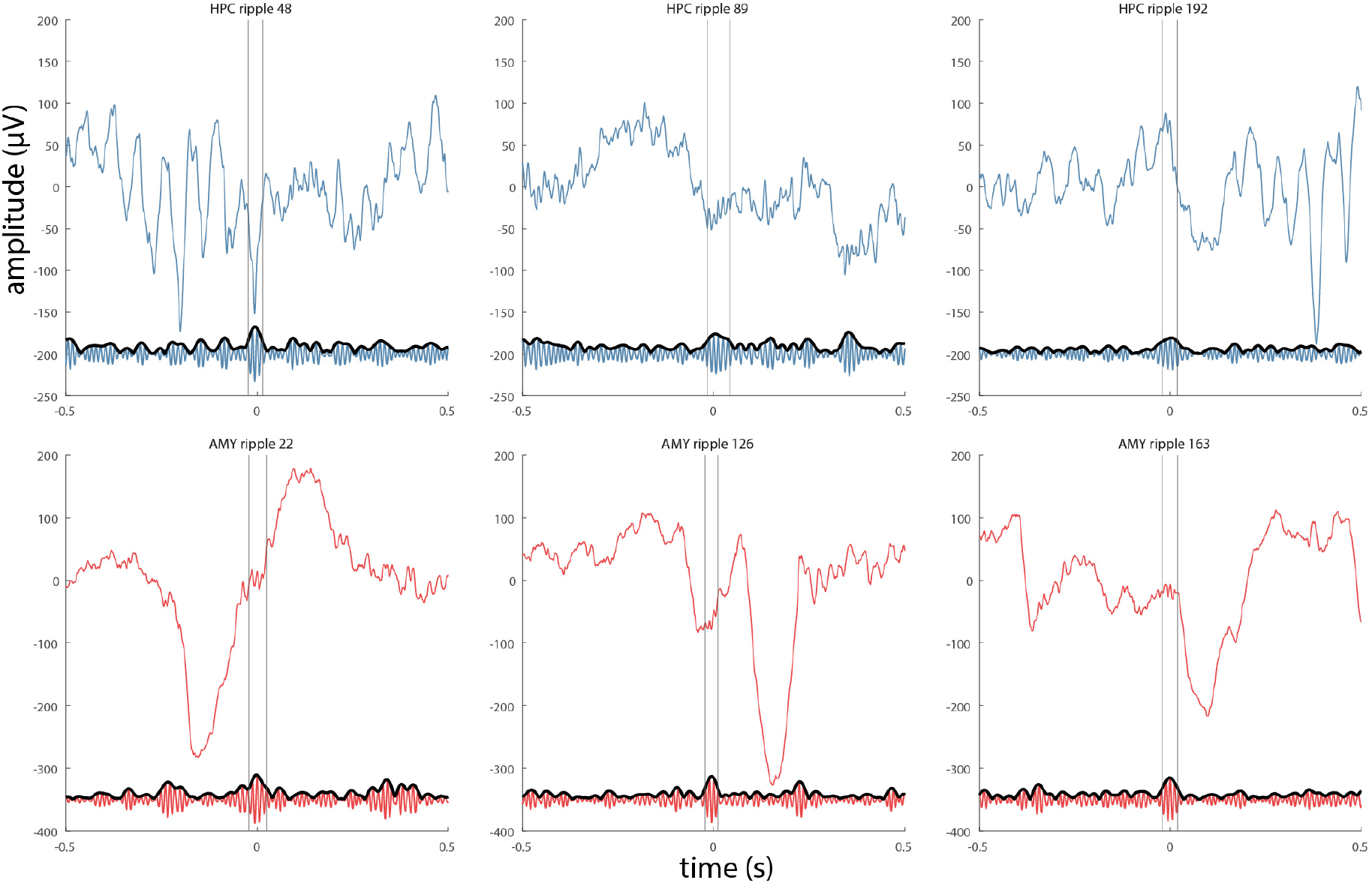
Examples of detected ripples for patient p4. Figure layout as in Figure 4. Note spindle rhythmicity in HPC (top-left and top-right panels).

**Supplementary Figure 4.**
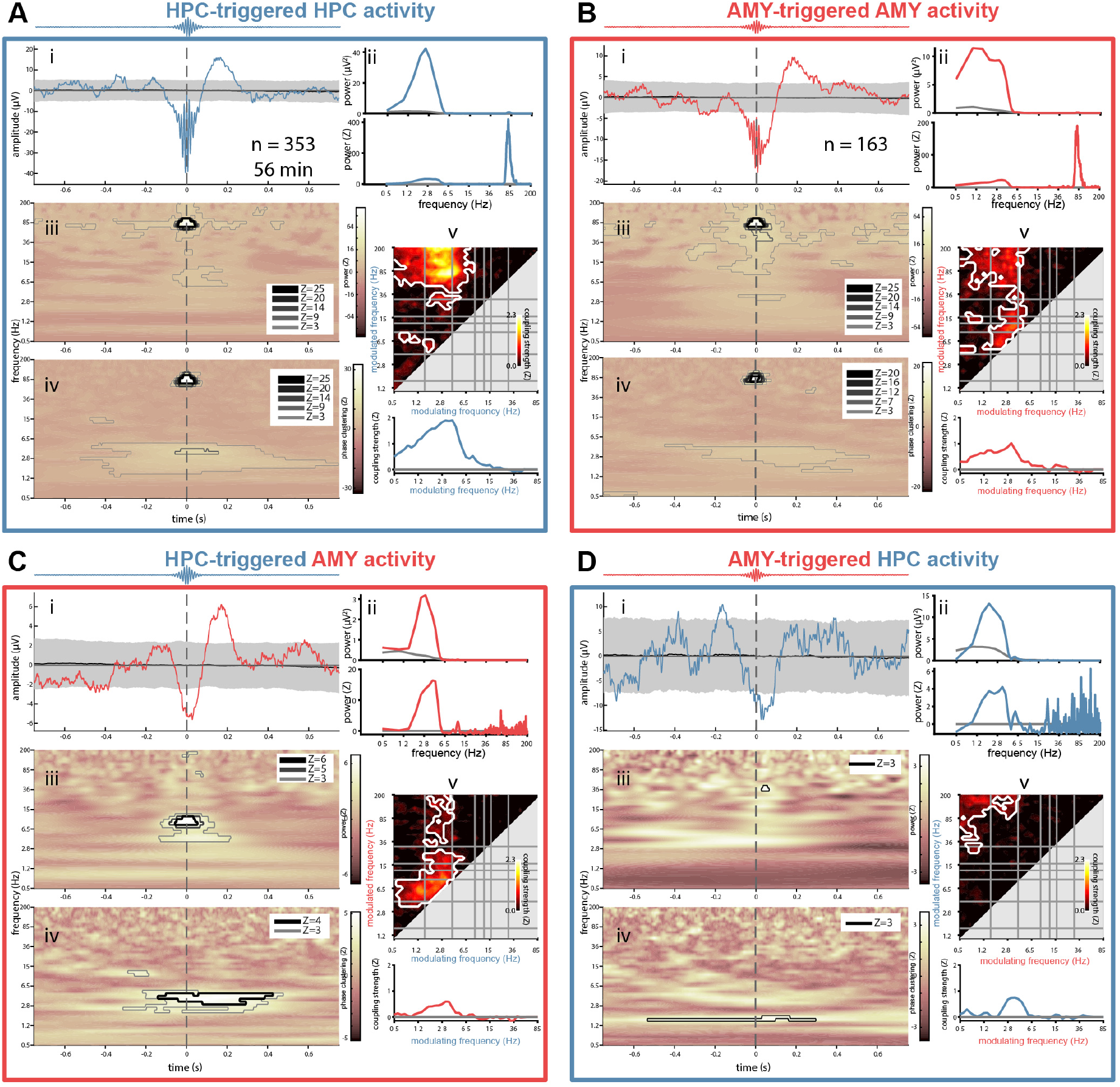
Local and interregional ripple-related dynamics in hippocampus and amygdala for patient p1. Panel layout identical to that of Figure 5.

**Supplementary Figure 5.**
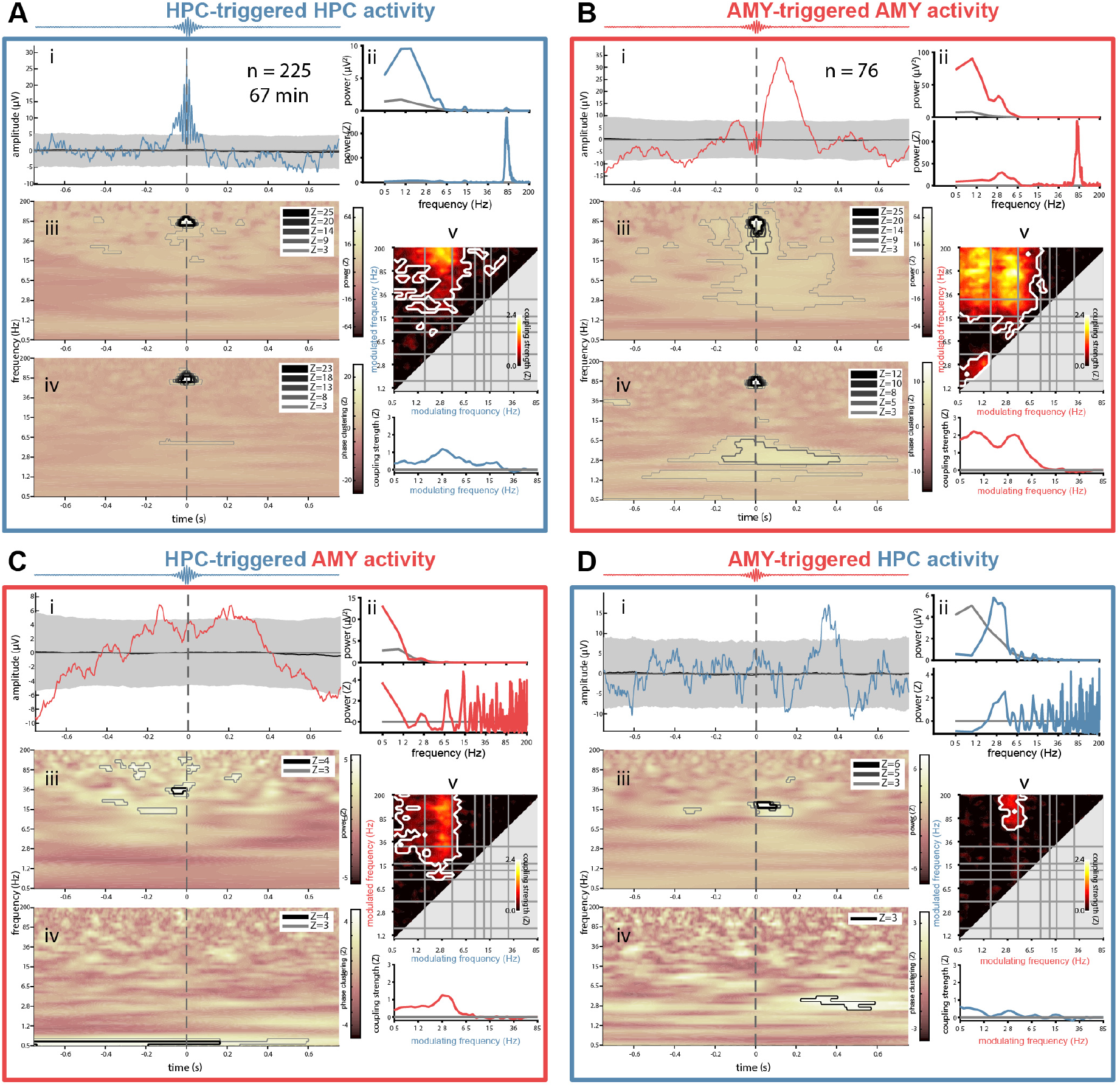
Local and interregional ripple-related dynamics in hippocampus and amygdala for patient p2. Panel layout identical to that of Figure 5.

**Supplementary Figure 6.**
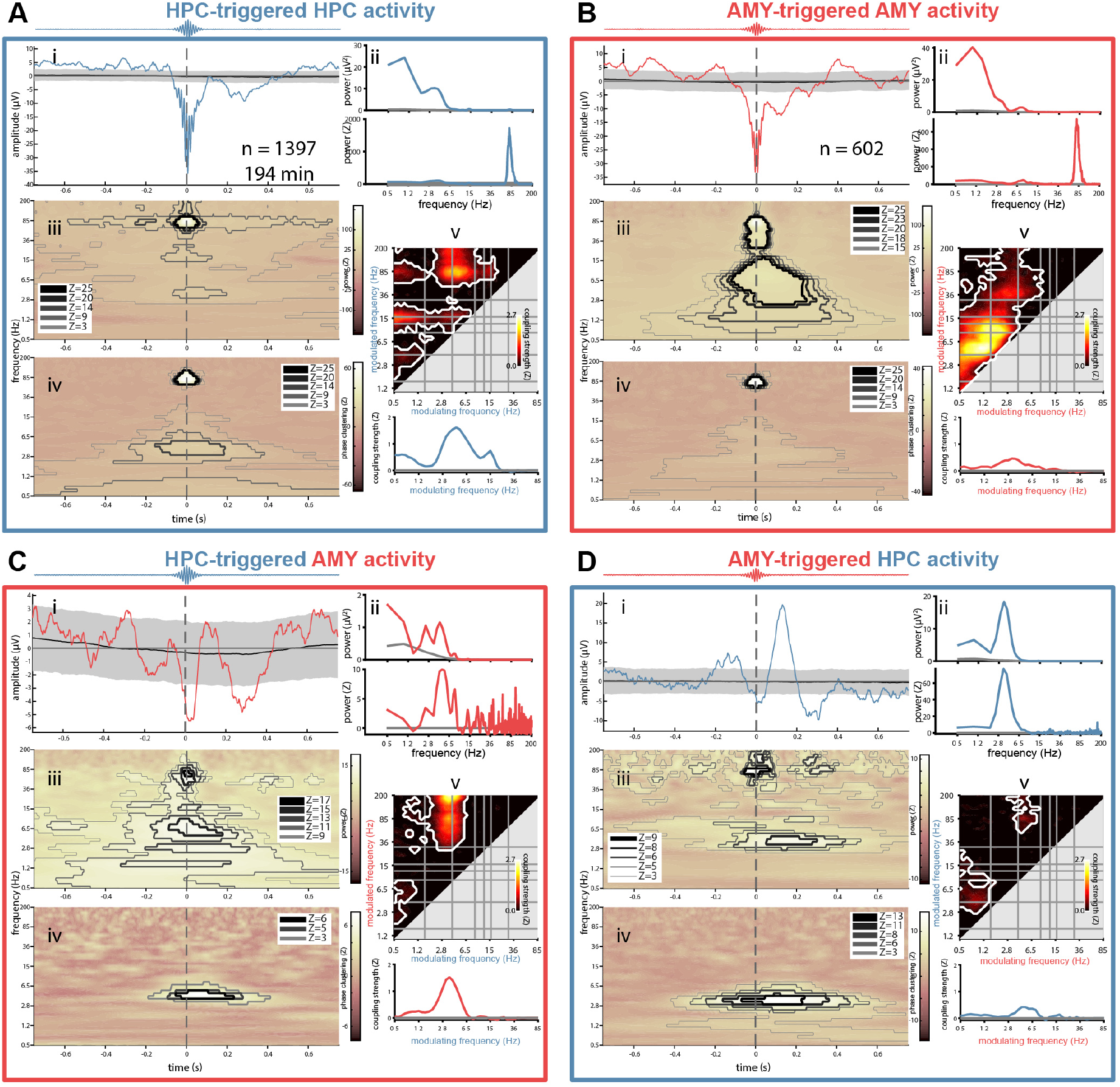
Local and interregional ripple-related dynamics in hippocampus and amygdala for patient p4. Panel layout identical to that of Figure 5.

## Notes

### Competing Interest Statement

The authors have declared no competing interest.

